# Experience shapes infants’ electrophysiological responses to faces from 3 to 9 months

**DOI:** 10.64898/2026.05.20.726644

**Authors:** Kira Ashton, Nicole Sugden, Wanze Xie, Fabiola Fernandez, Charisse B. Pickron, Margaret Moulson, Laurie Bayet

## Abstract

The types of faces that infants see impact their developing ability to engage with and individuate people from familiar and unfamiliar social groups, a phenomenon known as perceptual narrowing. However, the neural mechanisms that underlie infants’ processing of different faces as a function of experience remain poorly understood. To address this gap, the present study analyzes electroencephalography data collected while 3-month-olds (N=24), 6-month-olds (N=26), and 9-month-olds (N=18) viewed female and male faces of a familiar or unfamiliar social group. Infants’ neural responses to faces differed by group familiarity from 3 months of age, with increased responses to the more familiar face types in early components (P1, N290), and to the more unfamiliar face types in later components (P400, Nc). Face sex and group familiarity interacted to shape N290 and P400 amplitudes at 3- and 9-months. Specifically, N290 amplitudes were greater in response to female faces of a familiar group at 3 months, and to male faces of a familiar group at 9 months. In contrast, P400 amplitudes were greater in response to male faces of an unfamiliar group at 3 months old, and greatest in response to both female faces of a familiar group and to male faces of an unfamiliar group at 9 months. Source reconstruction of the Nc revealed greater reconstructed current density in response to faces of an unfamiliar social group across all ages. These findings contribute to a growing body of knowledge examining how perceptual experiences shape infants’ understanding of their social world.

---

The very first sources of social information that infants have access to include faces, and infants exhibit a visual preference for faces over non-face visual stimuli from birth (Mondloch et al., 1999; Viola Macchi et al., 2004). During the first year of life, infants’ face processing skills attune to the types of faces most present in their social environment, most notably, faces that match the race and gender of their primary caregiver (Bayet, 2022; Pickron & Bayet, 2026). Infants’ shifting behavioral and neural responses to faces of different races and genders shape their ability to perceptually encode and individuate diverse faces and are thought to contribute to the emergence of social biases and stereotypes (Quinn et al., 2020).

## Infants’ perception of faces of different races and genders

Infants’ perception of faces is shaped by the perceptual experiences and social interactions they encounter from an early age. In the first few months, infants develop a visual preference for faces of the same race as their primary caregiver (Kelly, Liu, et al., 2007). Around 9 months, infants’ visual preference for faces of a familiar race (own-race faces) begins to shift in the opposite direction, with an increasing looking preference for other-race faces (Fassbender et al., 2016; Kelly, Quinn, et al., 2007; Marquis & Sugden, 2019; Singarajah et al., 2017). At or around the same age, behavioral studies show a decrease in infants’ ability to distinguish between other-race faces, known as the "other-race effect" or perceptual narrowing (Kelly, Liu, et al., 2007; Sugden & Marquis, 2017). This phenomenon may be more common, or stronger, in infants raised in less racially diverse environments; conversely, some studies find no own-race bias in infants exposed to greater racial diversity (Pauker et al., 2016). Exposure to faces of a variety of races, ethnicities, and expressions fosters greater perceptual flexibility in infants, reflecting the adaptive nature of early perceptual development (Fort et al., 2024; Pauker et al., 2016).

Over the first year of life, infants’ face processing skills also become attuned to the gender of their primary caregiver (Liu et al., 2015; Quinn et al., 2008). Infants as young as 3-months old tend to visually prefer faces of the same gender as their primary caregiver, most often female (Quinn et al., 2002; Scherf & Scott, 2012). Importantly, while infants as a group tend to have more experience with female than male faces (due to unequal participation in early caregiving), infants’ individual experiences vary. Indeed, infants with a male primary caregiver show a visual preference for male rather than female faces (Quinn et al., 2002). Infants predominantly raised by a female primary caregiver appear less proficient at encoding or recognizing male than female faces, while infants with more balanced caregiving experiences do not (Ramsey-Rennels & Langlois, 2006; Rennels et al., 2017; Rennels & Kayl, 2020).

## Infant Event-Related Potentials associated with face processing

Event-related Potentials (ERPs) observed in infants in response to faces include the P1, N290, P400, and Nc (Conte et al., 2020). Of those, the N290 and the P400 are the closest analogs for the face-sensitive N170 component observed in adults (Leppänen et al., 2007) and are thought to reflect face-sensitive high-level visual processing and visual attention, respectively. Indeed, source reconstruction analyses (Michel et al., 2004) suggest that the infant N290 and the adult N170 both originate from the fusiform gyrus (Conte et al., 2020; Herrmann et al., 2005). Conversely, the P1 reflects low-level visual processing in both infants and adults (Tautvydaitė et al., 2022). The Nc is a frontal-central component associated with attention (Reynolds & Roth, 2018). Indeed, activity during the infant Nc time window is localized in the medial-anterior areas of the brain (Conte et al., 2020).

While the electrophysiological correlates of face processing are now well-established in infants, comparatively fewer studies have examined how these responses vary as a function of faces’ race or gender over the first year of life. N290 amplitudes are larger in response to female compared to male own-race faces in 7-month-old infants (Righi et al., 2014); at that age, the N290 also distinguishes novel from familiar female, but not male, own-race faces (Righi et al., 2014). Larger N290 amplitudes in responses to own-race avatar faces have also been reported at 9-months (Balas et al., 2011; Serafini & Pesciarelli, 2023; Vogel et al., 2012). The P400 also becomes sensitive to race during the first year, showing larger amplitudes in response to own than other-race female faces at 9-months, but not at 5-months (Grossmann, 2015; Vogel et al., 2012). Larger SSVEP responses to own-race compared to other-race female faces are also observed at 6- but not 9-months (Wallsinger et al., 2025). Finally, both familiarity and novelty can enhance the amplitude of the Nc, which is thought to reflect differential attention allocation (Reynolds & Roth, 2018) therefore, over the first year of life, Nc amplitudes may be expected to reflect the documented shift in attention allocation from faces of the most familiar groups to faces of the most novel racial groups (Marquis & Sugden, 2019). Overall, these findings suggest that the N290 and P400 become attuned to faces of the most familiar gender by 7-months, and to faces of the most familiar race by 9-months, with larger amplitudes observed in response to faces from the more familiar groups. However, the developmental trajectory of these experiential effects in the first year of life remains unclear; in addition, no study to date has examined the interactive effects of face race and gender on infant ERPs.

## Interactive effects of face-race and gender

Individual faces and social identities reflect the intersection of multiple dimensions, including but not limited to race and gender. Interactive effects of such factors shape the physical features of faces, as well as social experiences and biases, over and beyond their mere linear sum (Hester et al., 2020; Hudson et al., 2024). Less is known about how gender and race interact to shape infants’ face perception, although existing behavioral findings do suggest that some interactive effects emerge as early as 3 months. Infants between 3 and 6 months old demonstrate a visual preference for own-race, but not other-race female faces (Liu et al., 2015; Quinn et al., 2008). By at least 10 months, infants can categorize own-race faces by gender, yet they still partially fail to do so with other-race faces through at least 12-months of age (Damon et al., 2023). The “other-race effect” also emerges earlier for female than for male faces (Sugden & Marquis, 2017; Tham et al., 2015). These behavioral results suggest that the formation of face categories in infancy may already be driven by an interaction between the race and gender of faces, rather than solely by their additive effects.

## The present study

In summary, how infants’ looking behaviors in response to faces of different races and genders shift from 3 to 9-months of age has been well-described and reflects infants’ increasing attunement to their social and caregiving environment. However, the neural underpinnings of these developmental changes are less documented. In particular, the potential interactive effects of face-race and gender on infants’ neural responses to faces remain to be explored. To address these gaps, the current study sought to investigate the additive and interactive effects of face-race and face-gender on infants’ neural responses to faces at 3-, 6-, and 9-months of age.

Previous behavioral work shows that infants’ responses to faces’ race and gender shift over the first year of life, with some interactive effects of face-race and face-gender on infants’ visual preferences emerging as early as 3-4 months of age, and perceptual narrowing effects on face encoding or recognition emerging towards the end of the first year. Therefore, the first hypothesis was that significant differences in the amplitude of the attention-related Nc (and possibly P400; Guy et al., 2016) ERP component would be present as young as 3 months old in response to faces of different races and genders. Conversely, subtler differences in the N290 amplitude, reflecting expert face perception, were hypothesized to emerge later, by 9 months. The second hypothesis was that race and gender of the face stimuli would exert an age-dependent, interactive effect on the N290, P400, and Nc amplitudes. More specifically, it was hypothesized that 3-month-olds would show increased attention-related ERP amplitudes (i.e., Nc and possibly P400) to the most familiar faces (i.e., own-race female), shifting to increased attention-related ERP to the most novel faces (i.e., other-race male) by 9 months. Based on prior findings, at 9-months we additionally expected to observe larger N290 amplitudes in response to own-race female faces compared to other conditions (Balas et al., 2011; Righi et al., 2014), and larger P400 amplitudes in response to own-race compared to other-race female faces (Vogel et al., 2012). Finally, the third hypothesis was that electrical activity localized in brain regions previously associated with face processing in infants, such as the fusiform gyrus and occipital regions for the N290 and P400 time-windows, and frontal-central regions during the Nc time-window, would show effects of face race and gender similar to those evident in the corresponding ERPs.

## Methods

### Participants

Data were collected as part of a larger, longitudinal study of infant face processing. Informed consent was obtained from parents prior to the study sessions, and all research activities were approved by the [Redacted] University Research Ethics Board (now named [Redacted]). The final sample of included infants consisted of 24 3-month-olds (12 females, 98.25±11.80 days old), 26 6-month-olds (17 females, 187.70±10.17 days old), and 18 9-month-olds (11 females, 274.5±9.62days old), some of whom contributed longitudinal data at more than one time point. Data from additional infant visits (N=30 3-month-olds, N=23 6-month-olds, N=28 9-month-olds) were excluded due to the child declining to wear the cap, not completing a minimum of 10 total trials, or yielding fewer than 10 valid trials in each condition after preprocessing. Infants were invited to complete the study at multiple time points, with 3 of 50 unique included participants included at all three ages, 12 at two ages, and 35 at only one time point. For demographic information about the sample of included infants, see Table 1.

**Table 1.**
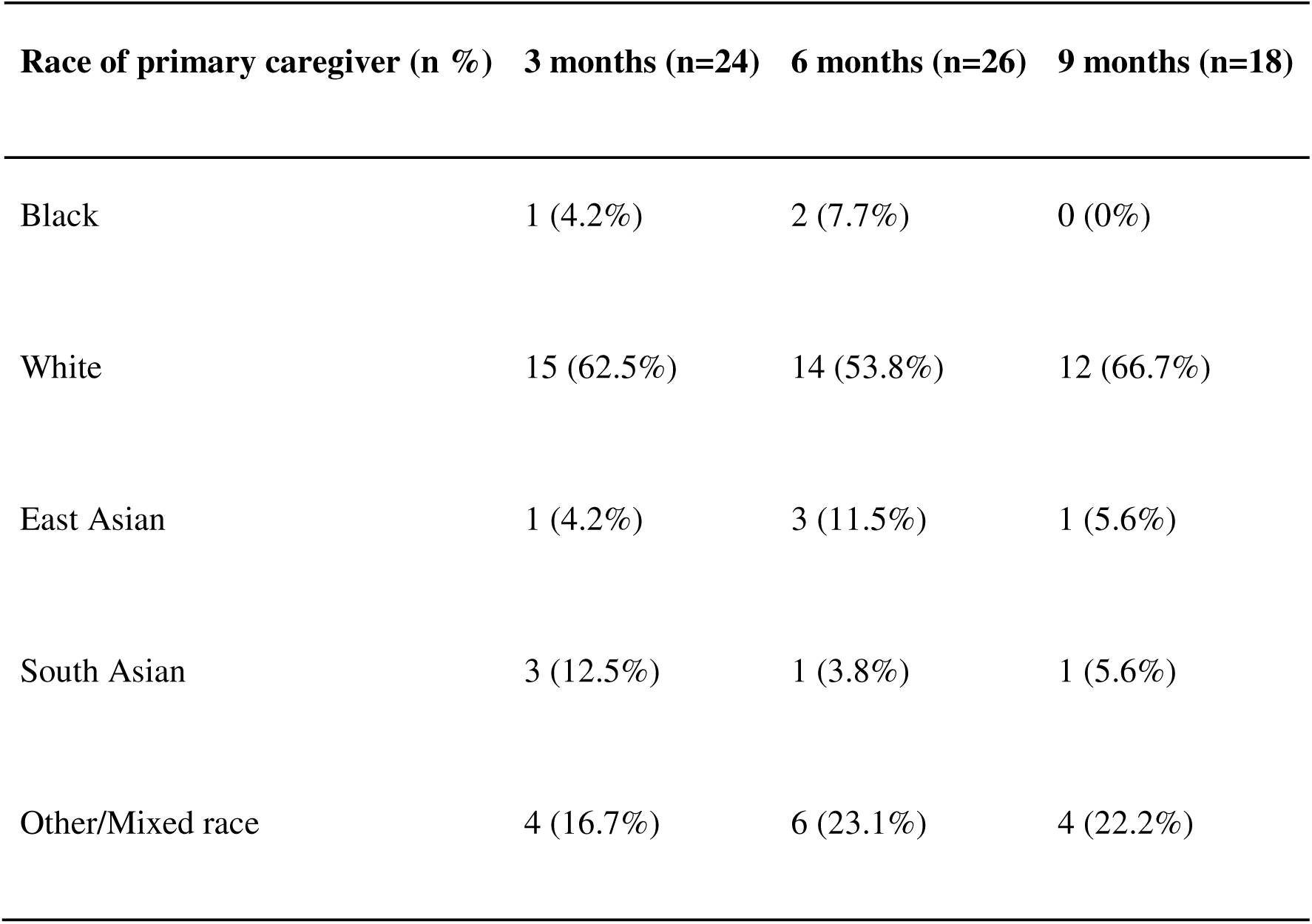
Participant demographics.

### Procedure

Infants passively viewed images of faces while seated on their parents’ lap, approximately 30 centimeters from a computer screen. Simultaneously, EEG was recorded using a high density 128-channel Hydrocel Geodesic Sensor Net (Electrical Geodesics Inc). EEG data was recorded using NetStation (EGI) with reference to Cz and sampled at 500 Hz. Infants were shown images of faces of male and female genders, either of the race of their primary caregiver (own-race) or of a different race than either of their parents (other race), resulting in 4 experimental conditions: other-race female faces, other-race male faces, own-race female faces, and own-race male faces. Infants saw pictures of 10 individual models of each condition, with one face of each type presented in each block from the Toronto Ethnically Diverse Face Database (Latif et al., 2025), presented at an approximate visual angle of 28.07 degrees, in random order for a maximum of up to 40 blocks (i.e., 160 trials). Racial and ethnic categories represented in the stimuli were selected to reflect the demographic makeup of the city where the data was collected. These categories included Black, White, East Asian, and South Asian. All models were slightly smiling. Faces were presented for 500 ms, followed by a 1 second blank screen and an attention-grabbing ball. The task was ended if the infant became too distracted or fussy, or the parent indicated that the infant was ready to stop.

### EEG Preprocessing

EEG data was pre-processed in NetStation. The continuous EEG signal was filtered at a high pass of 0.1 Hz, and a low pass of 30 Hz. The data was then epoched into 1100 ms segments, with a 100 ms baseline (pre-stimulus) period. Baseline correction was applied. Epochs were manually reviewed, and noisy channels were excluded on a trial-by-trial basis based on visual inspection by an experienced researcher. If more than 10% of channels were excluded on a given trial, that trial was rejected. Trials were also excluded if eye blinks, eye movements, or other artifacts were found. Spherical spline interpolation was used to interpolate noisy channels in remaining included trials. Lastly, data epochs were averaged within each condition and participant, and channels were re-referenced to their average.

### ERP Analysis

The P1 (90-170 ms), N290 (3 months: 250-390 ms; 6 and 9 months: 200-340 ms), P400 (3 months: 400-600 ms; 6 and 9 months: 350-600 ms), and Nc (300-700 ms) ERP components were considered for analysis (Guy et al., 2016). Time windows and channels of interest were selected based on work by Guy et. al. investigating cortical responses to faces in infants (Guy et al., 2016). Channels of interest for the P1, N290, and P400 included those situated nearest to visual and social processing areas of the cortex, specifically bilateral occipital (inion), bilateral occipito-temporal, and occipital midline channels (**Figure 1A**). For the Nc, channels of interest were those sensitive to areas of the cortex associated with executive function and attention, specifically the frontal midline, central midline, bilateral central channels (**Figure 1A**). For each participant and condition, the average amplitude of the P1, P400, and Nc over their respective time-windows and channels of interest was calculated (**Figure 2**). For the N290, peak amplitudes were computed during the N290 time-window for each participant and condition, then corrected by subtracting the peak P1 amplitude for that same participant and condition to minimize any potential residual effects of the P1 (Guy et al., 2022). The effects of age, ROI, face-race, and face-gender on ERP amplitudes were then assessed using linear mixed effect models using the lme4 package from R (Bates et. al., 2015), with a random intercept for participant ID.

**Figure 1.**
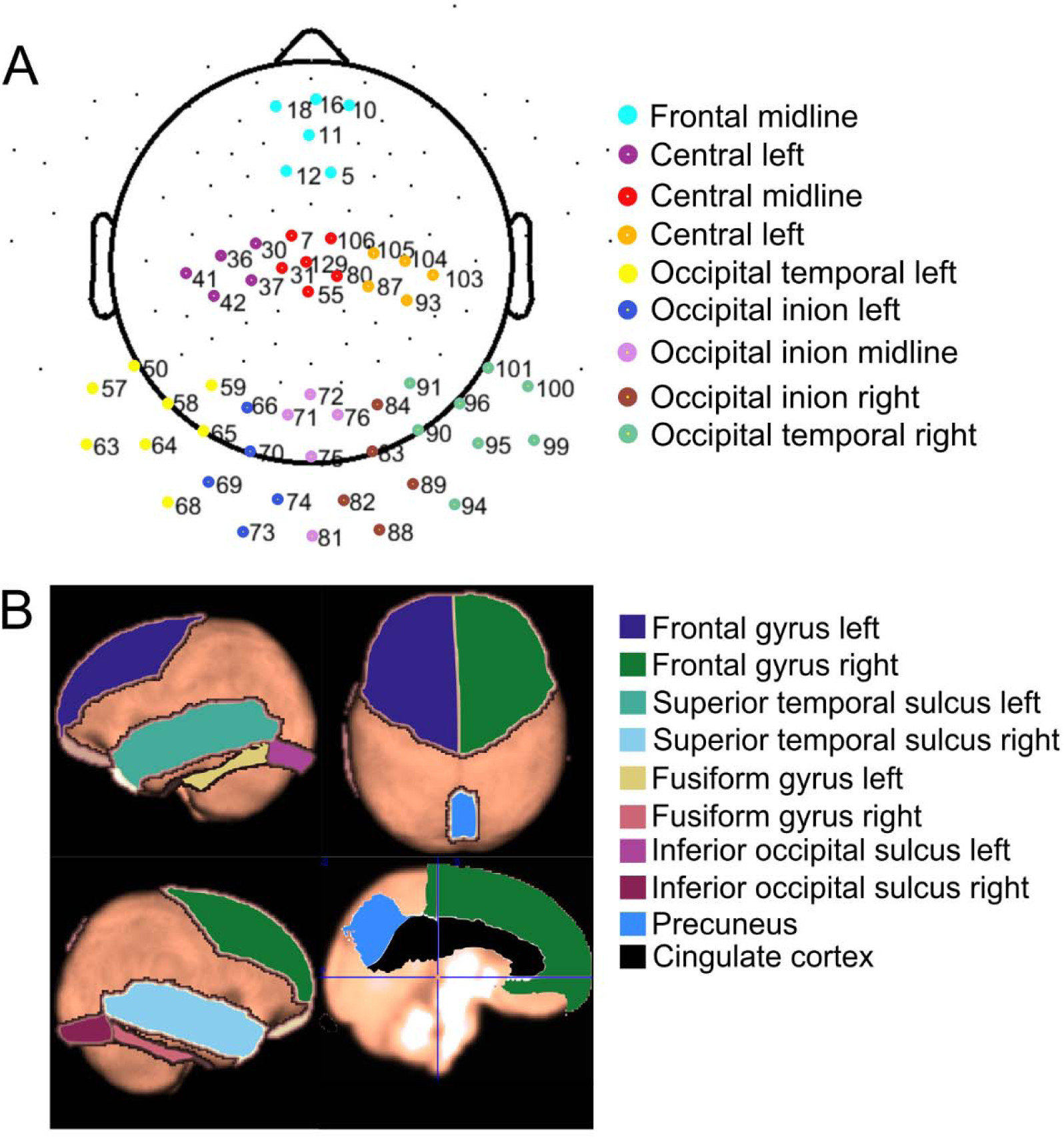
Regions of interest (ROIs) for (A) ERP analysis and (B) source reconstruction.

**Figure 2.**
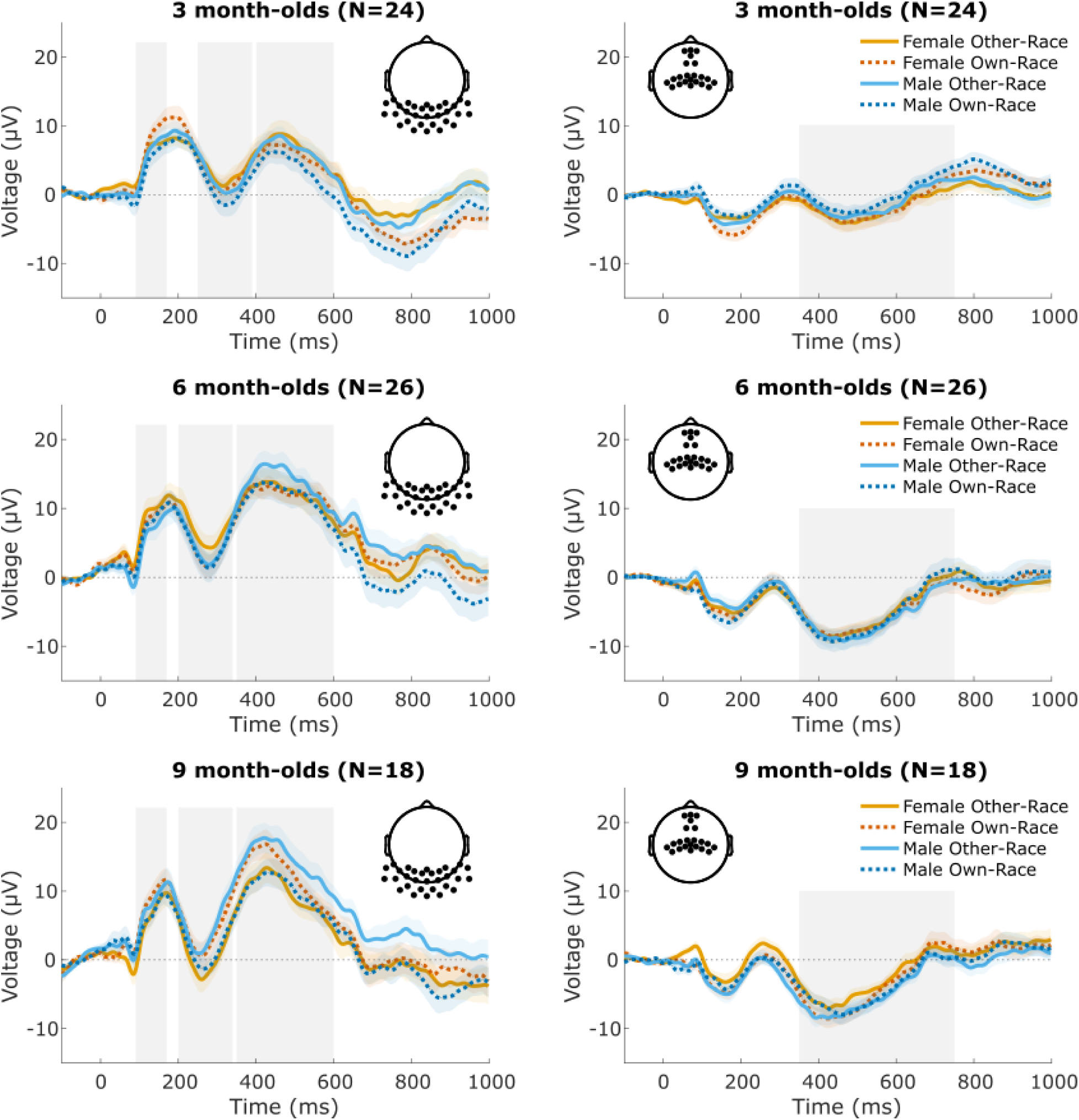
Average ERP waveforms and time-windows. Left: ERP amplitude averaged occipital and occipito-temporal ROIs for each age group, with P1, N290 and P400 time-windows highlighted. Right: ERP amplitude averaged over fronto-central ROIs for each age group, with the Nc time-window highlighted.

### Source localization

To identify specific cortical regions of interest that may contribute to the effects seen in ERP components, source localization was performed. Source localization first involved creating a head model using average template MRI images for each age group from the Neurodevelopmental MRI Database (Evans, 2006; Fillmore et al., 2015; Johnson, 2009; Richards, 2009; Richards et al., 2016; Richards & Xie, 2015; Sanchez et al., 2012). This database contains MRI data that has already been segmented into different tissue types (e.g., white matter, gray matter, CSF, skull, etc.). The head model was aligned with approximate electrode coordinates. Using the non-parametric eLORETA (exact low-resolution brain electromagnetic tomography) algorithm (Jatoi et al., 2014), a source model was constructed to estimate the current from any given source on a 5mm resolution grid for each of the 3 age groups. Using EEG data, the current density reconstruction (CDR) was calculated across sources, estimating the strength of activity at specific areas of the cortex, then averaged within ROIs and time-windows of interest. Cortical ROIs were selected based on prior work (Guy et al., 2016) to include areas of the cortex expected to be involved in face perception and visual attention in infants. For the activity occurring during the P1, N290, and P400 time windows, the ROIs were the bilateral fusiform gyri (FG), superior temporal sulci (STS, defined here as the conjunction of the superior and middle temporal gyri), and inferior occipital gyri (IOG; **Figure 1B**). Nc-related source activity was measured over the left and right frontal cortex (FC, defined as the conjunction of the superior and middle frontal gyri), precuneus (PC), and cingulate cortex (CC; **Figure 1B**). These regions were identified using the LPBA40 brain atlas (Rohlfing et. al., 2010). N290-related source activity was corrected for P1 activity by subtracting peak P1-related source activity. Finally, linear mixed effect regression models were used to examine the impact of age, ROI, face-race, and face-gender on the CDR, using lme4 in R (Bates et. al., 2015), with a random intercept for participant ID.

### Subset analyses

Some included infants viewed a stimulus condition (Black male faces, viewed either as own-race or as other-race faces) that differed from the others in terms of low-level visual properties (e.g., background color). Therefore, all analyses were repeated in a subset excluding trials where babies were viewing these less well-normed stimuli, to ensure that the reported findings in the full sample cannot be driven by such low-level visual differences (see **Supplementary Materials** for subset results).

## Results

### P1 Amplitude and related source activity

Analysis of the average P1 amplitude revealed significant main effects of ROI (*F*[4,1274]=131.71, *p<*.001; **Supplementary Figure 2**), age (*F*[1,1274]=5.54, *p*=.019), and face-gender (*F*[1,1274]=8.33, *p*=.004, **Figure 3**), with significantly greater P1 amplitudes in response to female than to male faces. P1 amplitudes were significantly higher in occipital inion ROIs than in the bilateral occipito-temporal ROIs (all *ps*<.001). Additionally, P1 amplitudes in the occipital inion midline were significantly greater than in the occipital inion left (*F*[1,493]=17.40, *p*<.001) and right (*F*[1,493]=5.99, *p*=.015) ROIs. Similar, significant main effects of face-gender and ROI on P1 amplitudes were found in subset analyses (**Supplementary Materials**) excluding trials with less well-normed stimuli, suggesting they are not driven by these stimuli. To investigate the possibility that luminance differences could account for the effect of face gender on P1 amplitudes, luminance values between male and female stimuli were compared and found to not significantly differ (independent *t*-test, *t*[76.62] = 0.53, *p* = 0.598).

**Figure 3.**
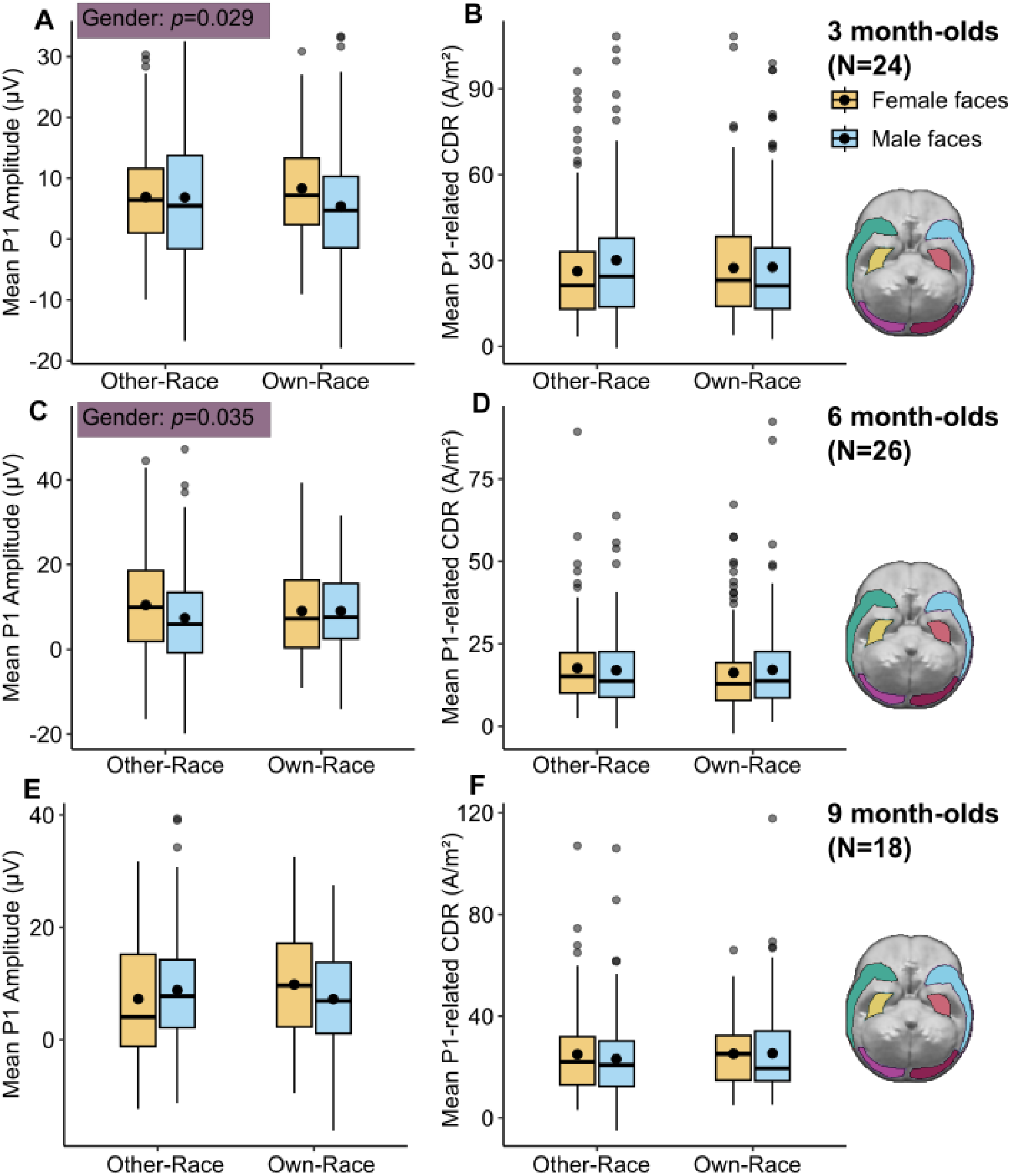
P1 activity by age and condition. Left: Average P1 amplitude over occipital and occipito-temporal ROIs. P1 amplitudes were significantly greater for female than for male faces at 3 and 6 months. Right: P1-related CDR activity averaged over occipital and occipito-temporal ROIs.

Analysis of the source activity (CDR) occurring over occipito-temporal ROIs during the P1 time-window revealed a significant interaction between ROI and age (*F*[5,1531] = 3.43, *p=*.004; **Supplementary Figure 3**). At 3 months, source activity was higher in the bilateral FG and IOG than in STS, and in the left compared to the right IOG (all *p*s < .05). At 6 months, source activity was higher in the left FG over right FG, bilateral IOG, and bilateral STS, in the right FG over the right IOG and bilateral STS, and in the left IOG over the left STS (all *p*s < .05). At 9 months, source activity was higher in the left FG over right FG, right IOG, and bilateral STS, in the right FG over right STS, and in the left IOG over right IOG and right STS (all *p*s < .05).

### N290 Amplitude and related source activity

P1-corrected N290 peak amplitudes showed a significant interaction between ROI and age (*F*[4,1274]=4.87, *p*<.001; **Supplementary Figure 2**) as well as an interaction between face-race, face-gender, and age (*F*[1,1274]=4.57, *p*=.033). These interactions held when restricting analyses to the better normed stimuli (see **Supplementary Materials**). At 3-months-old, N290 amplitudes were significantly greater (more negative) for female own-race faces as opposed to female other-race faces (*F*[1,215]=8.26, *p*=.005; **Figure 4A**), and a similar pattern was found in subset analyses (**Supplementary Materials**). At 6 months, N290 amplitudes in response to male own-race faces were significantly greater than to male other-race faces (*F*[1,232]=4.84, *p*=.029; **Figure 4B**); a numerically similar but only marginally significant pattern was found in subset analyses. There were no significant differences at 9 months. In addition, P1-corrected N290 amplitudes were significantly greater over occipital than bilateral occipito-temporal channels at all ages (*ps* <.001); at 3 months, amplitudes were also greater over occipito-temporal right than over occipito-temporal left channels (*F*[1,167]=8.95, *p*=.003).

**Figure 4.**
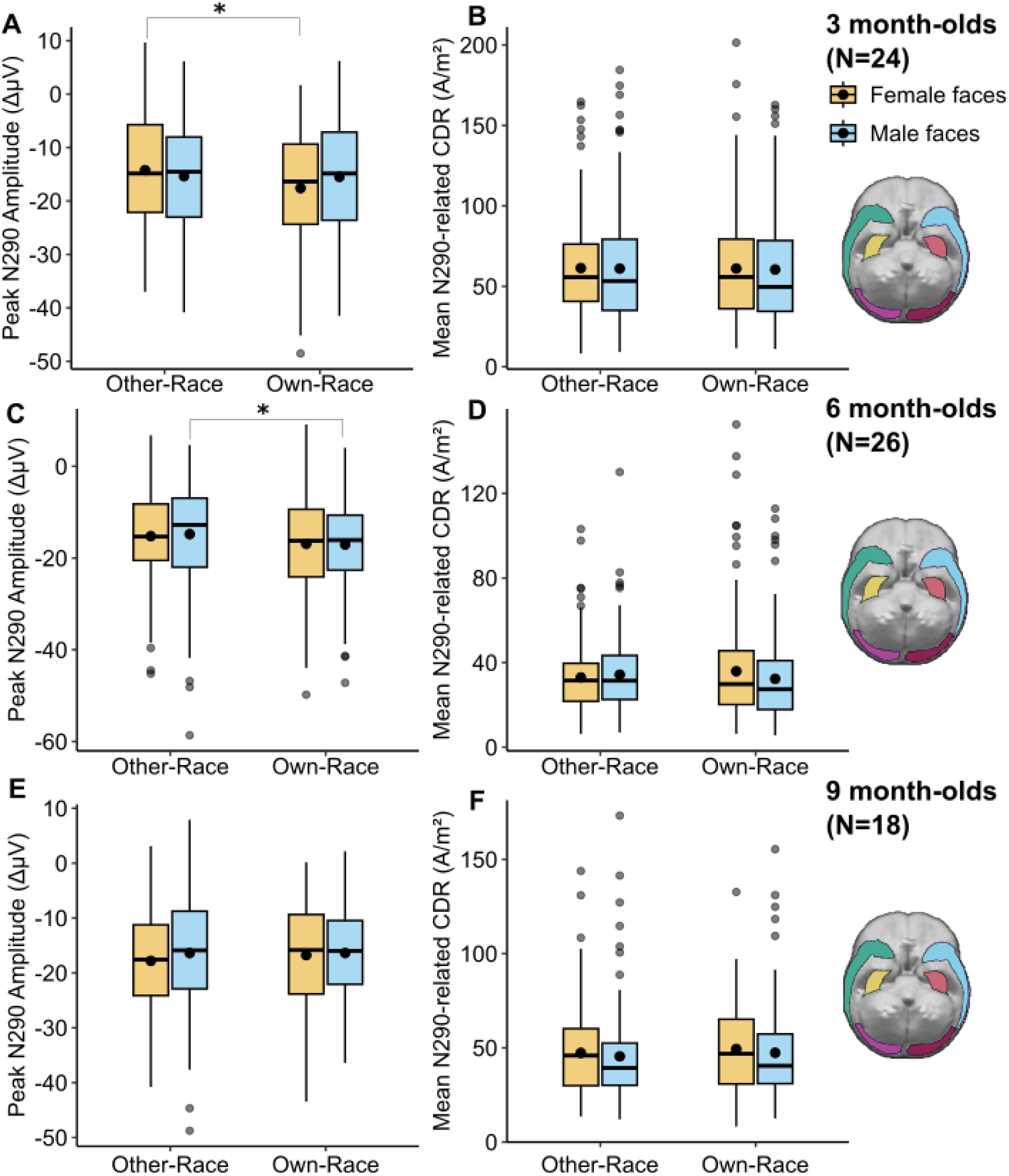
N290 activity by age and condition. Left: Peak N290 amplitude over occipital and occipito-temporal ROIs. At 3 months, N290 amplitudes were significantly greater in response to female own-race faces than female other-race faces. At 6 months old, N290 amplitudes were significantly greater in response to male own-race faces than male other-race faces. Right: N290-related CDR activity over occipital and occipito-temporal ROIs. No significant effects of face-race or face-gender.

A significant interaction between ROI and age (*F*[5,1531] =9.58, *p*<.001; **Supplementary Figure 3**) impacted source activity (CDR) in the N290 time window. At 3 months, CDR in the bilateral IOG and FG was significantly greater than in the bilateral STS (all *ps*<.001). At 6 and 9 months, CDR in the bilateral FG was significantly greater than in the bilateral STS and IOG (all *p*s<.001).

### P400 Amplitude and related source activity

The average P400 amplitude was significantly impacted by ROI (*F*[4,1274]=27.07, *p*<.001; **Supplementary Figure 2**), and a three-way interaction between face-race, face-gender, and age (*F*[1,1274]=3.99, *p*=.046). Specifically, P400 amplitudes were significantly greater over occipito-temporal rather than bilateral and midline occipital ROIs (*ps* <.001), At 3 months, P400 amplitudes were significantly greater for other-race than for own-race male faces (*F*[1,215]=5.80, *p*=.017; **Figure 5A**). At 9 months, P400 amplitudes were significantly greater in response to other-race male faces than to own-race male faces (*F*[1,161]=14.75, *p*<.001) or to other-race female faces (*F*[1,161]=11.06, *p*=.001) faces, and significantly greater in response to own-race female faces than to female other-race female faces (*F*[1,161]=4.06, *p*=.046) or to own-race male faces (*F*[1,161]=5.54, *p*=.020) faces (**Figure 5E**). In other words, at 9 months, P400 amplitudes were larger in response to of the most (female own-race) and least (male other-race) familiar types to infants. This three-way interaction (but not the interaction of age and ROI) remained significant when excluding trials during which infants saw non-normed stimuli (See **Supplementary Materials**).

**Figure 5.**
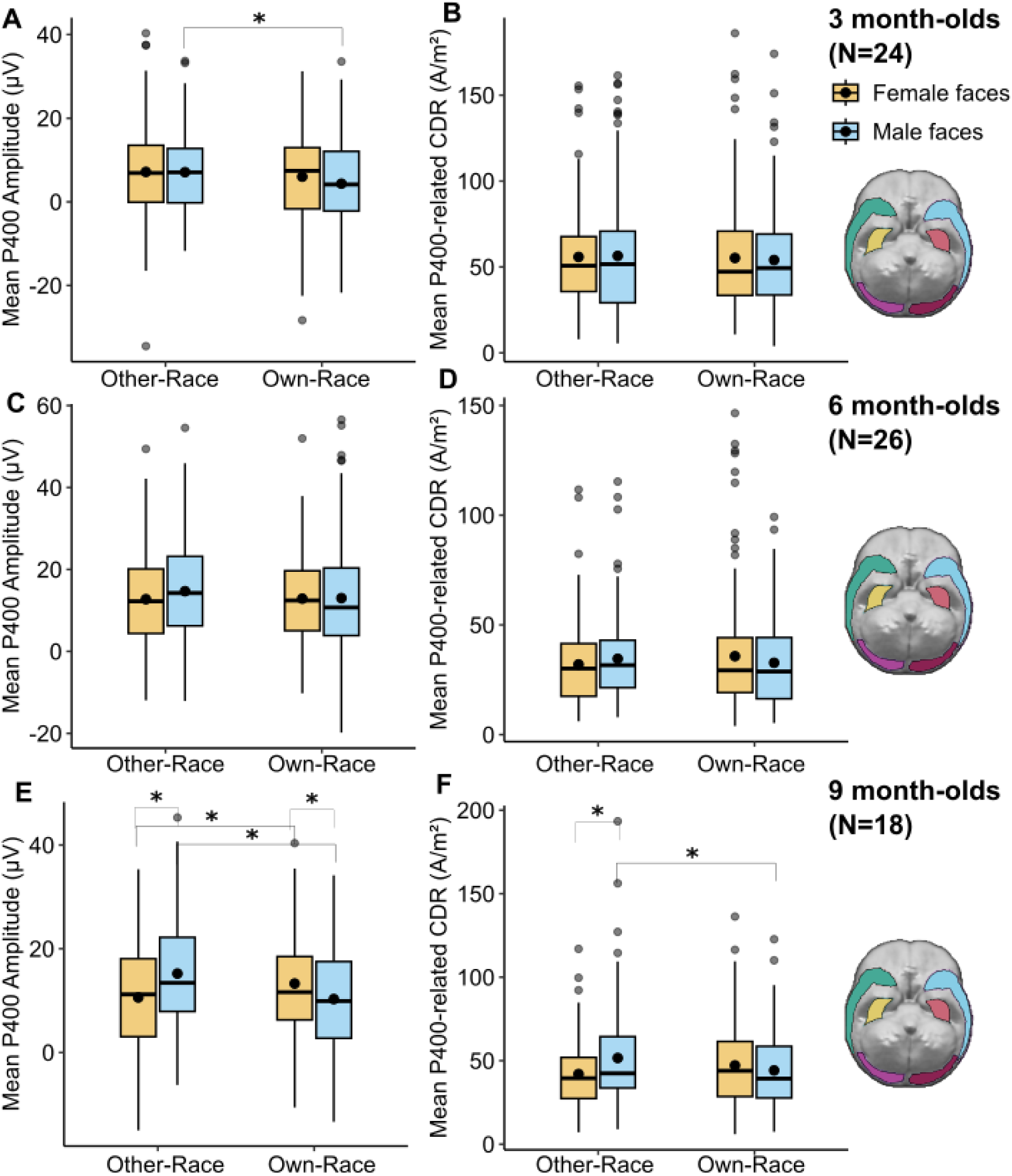
P400 activity by age and condition. Left: Average P400 amplitude over occipital and occipito-temporal ROIs. At 3 months, P400 amplitudes was significantly greater in response to male other-race faces than male own-race faces. At 6 months old there were no significant differences. At 9 months old, P400 amplitudes were significantly greater in response to male other-race faces than male own-race faces and female other-race faces, and in response to female own-race faces than to female other-race faces and male own-race faces. Right: P400-related CDR activity averaged over occipital and occipito-temporal ROIs. At 9 months, activity was greater in response to male other-race faces than to male own-race faces, or female other-race faces.

Corresponding source activity (CDR) in the P400 time-window was significantly impacted by an interaction between ROI and age (*F*[5,1531] =9.97, *p*<.001; **Supplementary Figure 3**), a significant interaction between face-race and face-gender (*F*[1,1531] =7.07, *p*=.008), and a marginally significant interaction between age, face-race, and face-gender (*F*[5,1531] =3.61, *p*=.058). The interaction between face-race and face-gender was only marginally significant when excluding trials during which infants saw non-normed stimuli, while the interaction between age and ROI remained significant (See **Supplementary Materials**). The interaction between face-race and face-gender was driven by larger CDRs in response to other-race male faces than to own-race male faces (*F*[1,739] =5.72, *p*=.017) or other-race female faces (*F*[1,743] =6.29, *p*=.012). Though the three-way interaction with age was not significant, this effect was only significant at 9-months (*F*[1,391]=10.09, *p*= .002, **Figure 5F**). In addition, at 3 months, CDRs were significantly greater in the left IOG and bilateral FG than in the bilateral STS (all *ps*<.001), and in the right IOG and left STS than in the right STS (*p*<.05). At 6 and 9 months, CDRs were significantly greater in the bilateral FG than STS and IOG, in the bilateral STS than in the right IOG, and in the left STS than in left IOG (all *p*s<.05); additionally, at 6 months, CDRs were slightly larger in the left than right FG (*F*[1,180] =4.03, *p*=.046).

### Nc Amplitude and related source activity

Average Nc amplitudes were impacted by infants’ age (*F*[1,1010]=23.06, *p*<.001). Nc amplitudes were significantly greater (more negative) at 6 months (*F*[1,758]=49.13, *p*<.001) and 9 months (*F*[1,634]=18.74, *p*<001) than at 3 months, and at 6 months than at 9 months (*F*[1,666]=8.14, *p*=.005).

Analysis of CDR values in frontocentral ROIs during the Nc time window revealed an interaction between ROI and age (*F*[5,1539] =7.66*,p<*.001; **Supplementary Figure 3**) and a main effect of face-race (*F*[1,1539] =7.74*, p*=.006; **Figure 6**). Specifically, frontocentral CDR values during the Nc time window were significantly stronger in response to other-race than to own-race faces. In addition, at 3 months, activity in the bilateral FC was significantly greater than in the bilateral CC and PC, and activity in the bilateral CC was significantly greater than in the PC (all *ps*<.01). At 6 months, activity in the bilateral PC was significantly greater than in the bilateral FC and CC (all *ps*<.05). At 9 months, activity was significantly greater in the left FC was significantly greater than in the bilateral CC, in the right FC than in the right CC, and in the right PC than in the right CC (all *ps*<.05).

**Figure 6.**
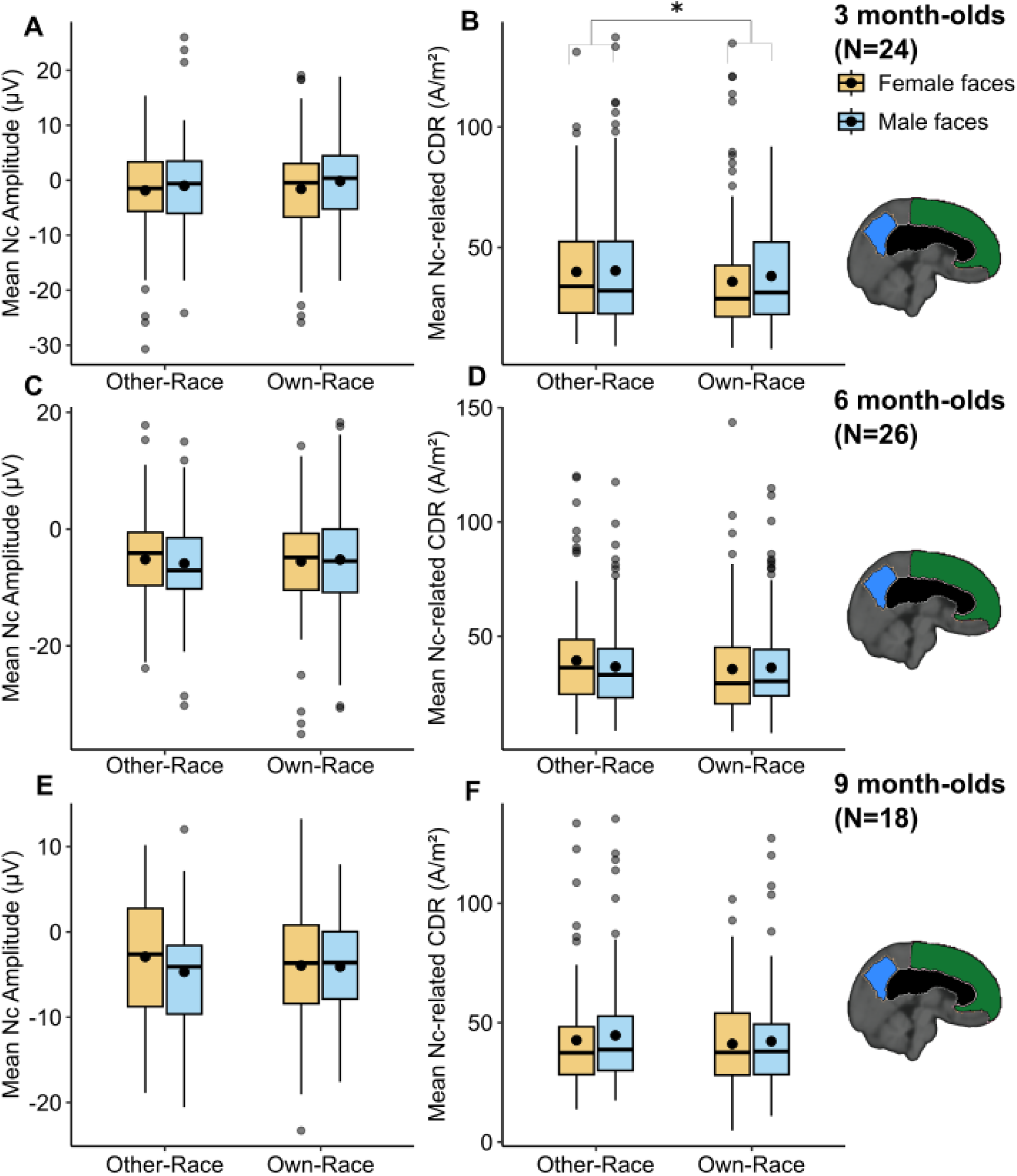
Nc activity by age and condition. Left: Nc amplitude over frontal/central ROIs during the Nc time window by face-race and face-gender conditions. No significant effects of face-race or face-gender. Right: Nc-related CDR activity averaged over frontocentral ROIs. At 3 months, activity was significantly greater in response to other-race faces.

These results were replicated in the subset analysis (see **Supplementary materials**).

## Discussion

The current study used EEG to examine additive and interactive effects of face-race and face-gender on infants’ neural processing of faces from 3 to 9 months of age, a developmental period during which infants exhibit experience-driven behavioral changes in their visual preference for and ability to individuate faces of different races or gender (i.e., perceptual narrowing). Face race and face gender interacted to impact N290 and P400 amplitudes in an age-dependent manner. In addition, cortical activity in frontocentral areas during the Nc time window was overall increased in response to other-race compared to own-race faces.

Our first hypothesis was that significant differences in the amplitude of the Nc component (indexing attention) in response to faces of different races and gender would be present as early as 3-months. while differences in the amplitude of the N290 component (reflecting expert face processing) would emerge later. Indeed, source analysis of the Nc revealed an overall pattern of elevated source activity in frontocentral regions in response to other-race faces compared to own-race faces across all age groups, suggestive of increased attention allocation to the more novel other-race faces as early as 3-months of age. However, contrary to our hypothesis, there were no effects of face race or gender on Nc amplitudes; age-dependent effects on N290 amplitudes were evident at 3- and 6-months, but not 9-months. Our second hypothesis was that face race and gender of the face stimuli would exert an age-dependent, interactive effect on the N290, P400, and Nc amplitudes. No such effect was expected on the P1, which is responsive to lower-level visual information. As predicted, N290 and P400 amplitudes differed by face race and gender in age-dependent manner. However, contrary to our hypothesis, no such effect was found on the Nc. In addition, P1 amplitudes were overall greater in response to female than to male faces. This unexpected effect may reflect a greater attunement of the earliest stages of face processing to female faces in infants (Rennels et al., 2017), or unmeasured low-level differences between female and male faces in the stimuli. To minimize the impact of such potential low-level effects on subsequent ERPs, P1-corrected N290 amplitudes were derived. Finally, our third hypothesis was that source activity in face processing regions of the cortex would show effects of face race and gender akin to those evident in the corresponding ERPs. Effects of face race and gender on source activity were similar to those found on channel ERPs for the P400 but diverged for other ERPs.

P1-corrected N290 amplitudes were stronger overall in response to own-race female than other-race female faces at 3-months, and in response to own-race male faces than other-race male faces at 6 months. These results align with prior work generally reporting larger N290 amplitudes in response to faces of the more familiar group in infants, such as in response to human compared to monkey faces at 6-months (de Haan et al., 2002) in response to female compared to male own-race faces at 7-months (Righi et al., 2014), and in response to own-race compared to other-race avatar faces at 9-months (Balas et al., 2011). No effects of face race or gender were found on N290 amplitudes at 9-months, in line with a previous report of similar N290 amplitudes in response to own- and other-race female faces at this age (Vogel et al., 2012). The current findings additionally suggest that the N290 may already be sensitive to the familiarity of racial characteristics by 3-months of age, with potentially more expert face processing of own- compared to other-race female faces at 3 months, and of own- compared to other-race male faces at 6 months, consistent with an ongoing specialization for the more frequently encountered types of faces.

An age-dependent interactive effect of face-race and gender also shaped P400 amplitudes, which are thought to reflect the allocation of attention to faces specifically (Conte et al., 2020). At 3-months, P400 amplitudes were larger in response to other-race male compared to own-race male faces. However, behaviorally, 3-month-olds tend to visually prefer own-race faces over other-race faces (Kelly, Quinn, et al., 2007), and female over male own-race faces, with no visual preference between other-race female and other-race male faces (Quinn et al., 2008). Therefore, the present ERP findings at 3-months contrast with previous behavioral findings in this age group. However, previous work has reported either no visual preference for own-race faces or a small numerical preference for other-race faces in the current cohort of infants (Sugden, 2016); the current ERP results align with these findings. Because infants in the current cohort were raised in a racially diverse Canadian city (Redacted), it is possible that their exposure to other-race faces sufficed to orient their attention towards other-race and own-race faces more equally. At 9-months, P400 amplitudes were larger in response to faces of the most familiar (female own-race) and most novel (male other-race) groups. This interactive effect of face-race and face-gender on P400 amplitudes was replicated in subset analyses, and a similar effect was evident in P400-related cortical activity. A previous ERP study, which used only female faces, also reported similar P400 amplitudes to own-race and other-race females faces at 5-months, but larger P400 amplitudes to own-race over other-race female faces at 9-months (Vogel et al., 2012). The current results partly align with those earlier findings, with no difference in P400 amplitude by face race or gender at 6 months of age, and increased P400 amplitudes to own-race versus other-race female faces at 9-months. Beyond perceptual familiarity or novelty, faces from the most familiar group might evoke safety and valuable social information to 9-month-old infants, while faces from the most unfamiliar group might evoke a potential threat. Indeed, older infants associate other-race faces with sad music, and own-race faces with happy music (Xiao et al., 2018). Regardless of affective valuation, the current results suggest that differential P400 amplitudes by face-race and gender emerge by 3-months, reorganize to show no difference around 6-months, then re-emerge by 9-months as experience further shapes face processing networks.

This study has several limitations. First, there was no measure of the racial or ethnic diversity of each individual infant’s social environment. The current study was conducted in [Redacted], a highly diverse city where 55.7% of residents identify as belonging to a non-white racial group ([Redacted] Census, 2021). However, infant head-camera data at the study site found that young infants’ visual experience with faces was nonetheless dominated by the faces of their primary caregiver, and therefore, by female and “own-race” faces (Sugden et al., 2014; Sugden & Moulson, 2019). Relatedly, and while early caregiving remains heavily female-biased overall (Minkin & Horowitz, 2023), the extent to which individual infants in the current sample had a female primary caregiver is unknown. Individual differences in the gender roles of a particular family would be expected to alter the observed differences in responses to female and male faces (Quinn et al., 2002). A second limitation is that the number of infants with included data at more than one time-point was too small to carry out longitudinal analyses. Future studies may address these gaps by relating granular individual experiences to differential neural responses, ideally longitudinally. Third, one stimulus condition was less standardized than the others, potentially introducing low-level confounds. To minimize this risk, all analyses were repeated with the affected trials excluded, and N290 amplitudes were corrected for P1 effects. Future work may leverage more recent, diverse face stimuli sets, e.g., the Chicago Face Database (Ma et al., 2021) to replicate the current findings with more fully normed stimuli.

The present study adds to our understanding of the effects of experience on infants’ neural processing of faces. Face race shaped infants’ neural responses to faces as early as 3-months of age, with generally heightened responses to the more familiar face types in early, perceptual components (P1, N290), and to the more novel face types in later, attention-related component (P400, Nc). Face race and face gender interacted to shape expert face processing (as indexed by N290 amplitudes) and attention allocation to faces (as indexed by the P400 amplitudes) as early as 3-months of age. In particular, at 9-months, P400 amplitudes were larger in response to both the most familiar (female own-race) and unfamiliar (male other-race) face types. These results demonstrate the need to employ diverse face stimuli that differ along intersecting dimensions of identity and include male faces, to better capture infants’ emerging understanding of the social world. Early differences in perceptual attunement and attention allocation towards individuals from different groups are thought to contribute to the formation of face categories and early perceptual-to-social biases (Markant & Scott, 2018). Indeed, attention allocation is one mechanism by which early behavioral interventions are thought to overcome perceptual narrowing (Hills & Lewis, 2006; Markant et al., 2016; Zhou et al., 2014), with potential implications for reducing implicit social biases (Lee et al., 2017). The emergence of neural tuning and differential processing of faces from familiar and unfamiliar groups by 3 months suggest that such interventions may be timed from early infancy.

## Funding statement

LB acknowledges support from a Young Investigator Award from the Brain & Behavior Research Foundation, a Karen Toffler Charitable Foundation award, and a CAS Collaborative Pilot grant from American University. MM acknowledges support from a Natural Sciences and Engineering Research Council (NSERC) Discovery Grant (#RGPIN-2019-05548).

## Conflict of interest disclosure

None to report.

## Ethics approval statement

Informed consent was obtained from parents prior to the study sessions, and all research activities were approved by the Ryerson University Research Ethics Board (now named Toronto Metropolitan University).

## Acknowledgments

We thank the infants and families who participated in this research.

## Notes

### Competing Interest Statement

The authors have declared no competing interest.

